# Integrin-activating Yersinia protein Invasin sustains long-term expansion of primary epithelial cells as 2D organoid sheets

**DOI:** 10.1101/2024.09.03.610788

**Authors:** Joost J.A.P.M. Wijnakker, Gijs J. F. van Son, Daniel Krueger, Willine van de Wetering, Carmen Lopez-Iglesias, Robin Schreurs, Fenna van Rijt, Sangho Lim, Lin Lin, Peter J. Peters, Ralph R. Isberg, Claudia Yanda, Wim de Lau, Hans Clevers

## Abstract

Matrigel/BME, a basement membrane-like preparation, supports long-term growth of epithelial 3D organoids from adult stem cells (ASC)^1,2^. Here, we show that interaction between Matrigel’s major component Laminin111 with epithelial α6β1-integrin is crucial for this process. The outer membrane protein Invasin of *Yersinia* is known to activate multiple integrin-β1 complexes, including integrin-α6β1. A C-terminal integrin-binding fragment of Invasin, coated on culture plates, mediated gut epithelial cell adhesion. Addition of organoid growth factors allowed multi-passage expansion in 2D. Polarization, junction formation and generation of enterocytes, goblet cells, Paneth cells, and enteroendocrine cells was stable over time. Sustained expansion of other human-, mouse-, and even snake epithelia was accomplished under comparable conditions. The 2D ‘organoid sheet’ format holds advantages over the 3D ‘in gel’ format in terms of imaging, accessibility of basal and apical domains and automation for high throughput screening. Invasin represents a fully defined, affordable, versatile, and animal-free complement to Matrigel/BME.

## Introduction

Epithelial cells *in vivo* require a close interaction with the basal lamina to establish correct apical-basal polarity and to prevent anoikis^3,4^. The basal lamina is composed of multiple ECM molecules, including collagens, Arg-Gly-Asp (RGD)-containing proteins and laminins^5,6^. Laminin molecules are large (800-900 kDa) multimers consisting of α, β and γ chains^7^. Epithelial cells interact with ECM components through different combinations of basally expressed αβ-heterodimeric integrin receptors (Suppl. Fig. 1)^8^. The α-chains each interact with specific ECM components while the common β-chain mediates downstream signals that control actin cytoskeletal rearrangements, polarity and cell survival^9,10^. Thus, integrin α5β1 recognizes RGD-containing ECM-proteins such as fibronectin, whereas α6β1 binds the embryonic laminin111 isoform (composed of laminin α1, β1 and γ1 chains)^11,12^(Suppl. Fig.1).

For *in vitro* culturing of primary epithelial cells, activation of integrin complexes is crucially dependent on Matrigel^13^. Matrigel^®^ and BME^®^ are the commercial names of an extract of Engelbreth-Holm-Swarm sarcoma tumor cells, grown in the abdomen of mice^14^. Matrigel/BME mainly consists of the embryonic laminin111 and additionally contains collagen IV, fibronectin and multiple minor components^15,16^. (Of note, integrin α3β1 binds adult-stage laminins that are not present in Matrigel/BME). Matrigel/BME is soluble at 4°C and gel-like at 37°C. Epithelial cells embedded in Matrigel/BME form polarized 3D mini-structures, also known as organoids^4,17^.

We have previously shown that the addition of a niche-mimicking cocktail of growth factors to single gut epithelial stem cells embedded in Matrigel/BME allows the long-term expansion of organoids retaining many structural and functional characteristics of the original tissue^1,2^. Nowadays, most primary epithelia can be grown under such defined 3D conditions^18^. Many studies have proposed alternatives to Matrigel/BME, for instance by functionalizing synthetic hydrogels with fragments of ECM proteins^19–21^, by using purified ECM proteins such as collagens^22^ or by extracting ECM material from post-mortem animal-or human organs^15,23,24^. However, often these hydrogels lack the key signalling molecule (laminin) present in Matrigel/BME to obtain its full protentional to support organoid growth. It appears fair to state that no material has been described to date that fully replaces Matrigel/BME. In the current studied, we sought to address if bacterial integrin-activating proteins can serve as alternatives to Matrigel/BME.

## Results

We first aimed to identify the integrin receptor(s) of colon epithelial cells that are required for Matrigel/BME to induce cell polarity, survival, and growth. By mRNA sequencing, we deduced that the integrins α6, αV and their partner chain β1^8^ represent the dominantly expressed integrin chains on these cells with the relevant affinity for Matrigel/BME components (Fig. 1A). We then tested the individual effects of blocking these three integrins. The function of integrin β1 was inhibited by adding the allosterically blocking antibody AIIB2 to the culture medium, which resulted in the rapid demise of the organoids (Fig. 1B,C). The *ITGA6* and *ITGAV* genes were mutated by CRISPR-Cas9 base-editing^25^(Suppl. Fig. 2). As these integrins expectedly generate essential growth signals, we added Rho-kinase inhibitor (Y-27) to the mutant organoid cultures to circumvent the loss of these signals^26^. Indeed, mutant organoid lines could readily be expanded in presence of Y-27. We re-plated equal numbers of wildtype (wt) and mutant organoid cells under standard organoid growth conditions in Matrigel/BME. This revealed that growth of these epithelial organoids fully depended on α6β1 integrin, and to a lesser extent on αVβ1 integrin (figure 1B). Of note, the α6β1 integrin receptor is the only available integrin receptor on colon organoids that recognizes the LN-111 protein^12^, the embryonic laminin that is abundantly present in BME and Matrigel.

**Fig. 1:**
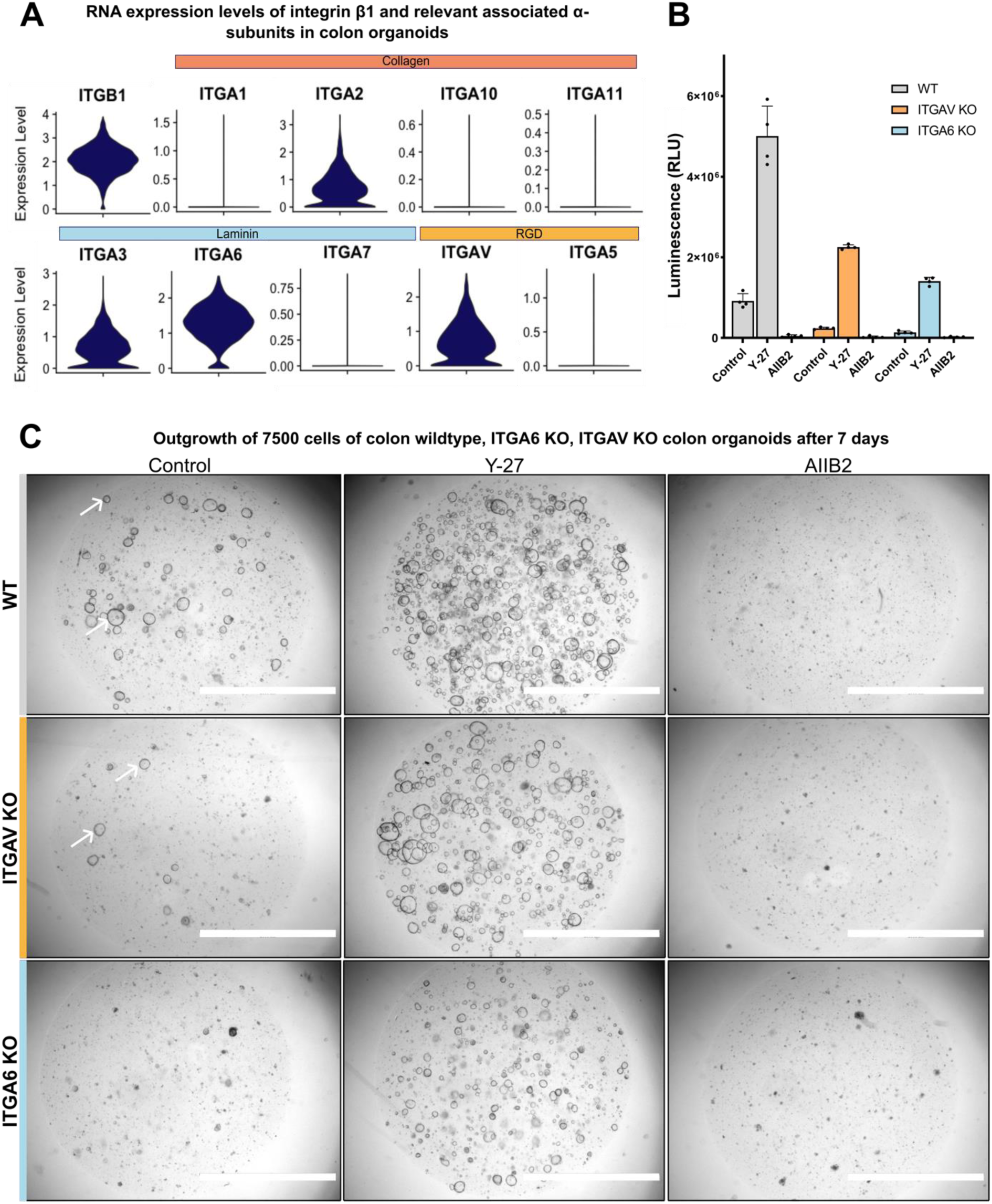
α6β1 is essential for colon organoid growth in Matrigel/BME. **A)** Single cell RNA sequence data of colon organoids (from^27^), indicating the expression levels of α-integrin subunits, selected for their interact with collagens, laminins or RGD proteins. The presented α-integrins all dimerize with integrin β1. Their ligands are indicated in the bars above the violin plots. **B/C** 7500 single cells of wildtype (WT), ITGA6 and ITGAV knockout (KO) colon organoids were cultured in BME^®^ hydrogels in standard medium, with Rho-kinase inhibitor (Y-27) or the β1-integrin allosterically inhibiting antibody AIIB2. Cultures were cultured for 7 days. **B)** Quantification of viable cells using ATP-sensitive luminescent assay (CellTiter-GLO). Mean ± SD, n=4. **C)** Bright-field images of the cultures. Scale bar: 2mm.

Our first attempt to replace Matrigel/BME by a bacterial integrin-activating protein was focused on the CagL protein of *Helicobacter pylori*^27^ and failed (not shown). We then turned to the enteropathogenic *Yersinia*. This Gram-negative bacterium employs its surface-expressed Invasin protein to enter its host through a rare intestinal epithelial cell type, the microfold (M) cell which overlies lymph node-like structures called Peyer’s patches^28–30^. M-cells uniquely express integrins on their luminal (rather than basal) surface^29,31^. *Yersinia* can reportedly bind and activate integrin-α3β1, -α4β1, -α5β1, -α6β1 and -αVβ1^32,33^ (Suppl. Fig. 1), through the small 25 kD C-terminal domain of its outer membrane protein Invasin (Inv192). This interaction is well studied from the perspective of allowing *Yersinia* to invade through the luminal surface of M-cells^34^. The specific interaction of Invasin with integrin complexes has previously been studied almost exclusively in immune cells^35^. At least 17 Yersinia species have been described. Among those, three are pathogenic in man: *Y. pestis, Y. pseudotuberculosis* and *Y. enterocolitica*^36^. Most *Yersinia* species harbour homologs of the Invasin (*Inv)* gene^37^. The two enteropathogenic *Y.pseudotuberculosis* and Y.*enterocolitica,* distantly related to each other^37^, both invade M-cells using their Invasin protein^29,30^. Figure 2A gives an alignment of the well-defined integrin-activating C-terminus (Inv192) of these two Invasin homologs. Of note, *Y. pestis* is unable to invade epithelial cells due to an insertion of an unrelated nucleotide sequence into the central region of its -otherwise intact-*Inv* gene^38^.

**Fig. 2:**
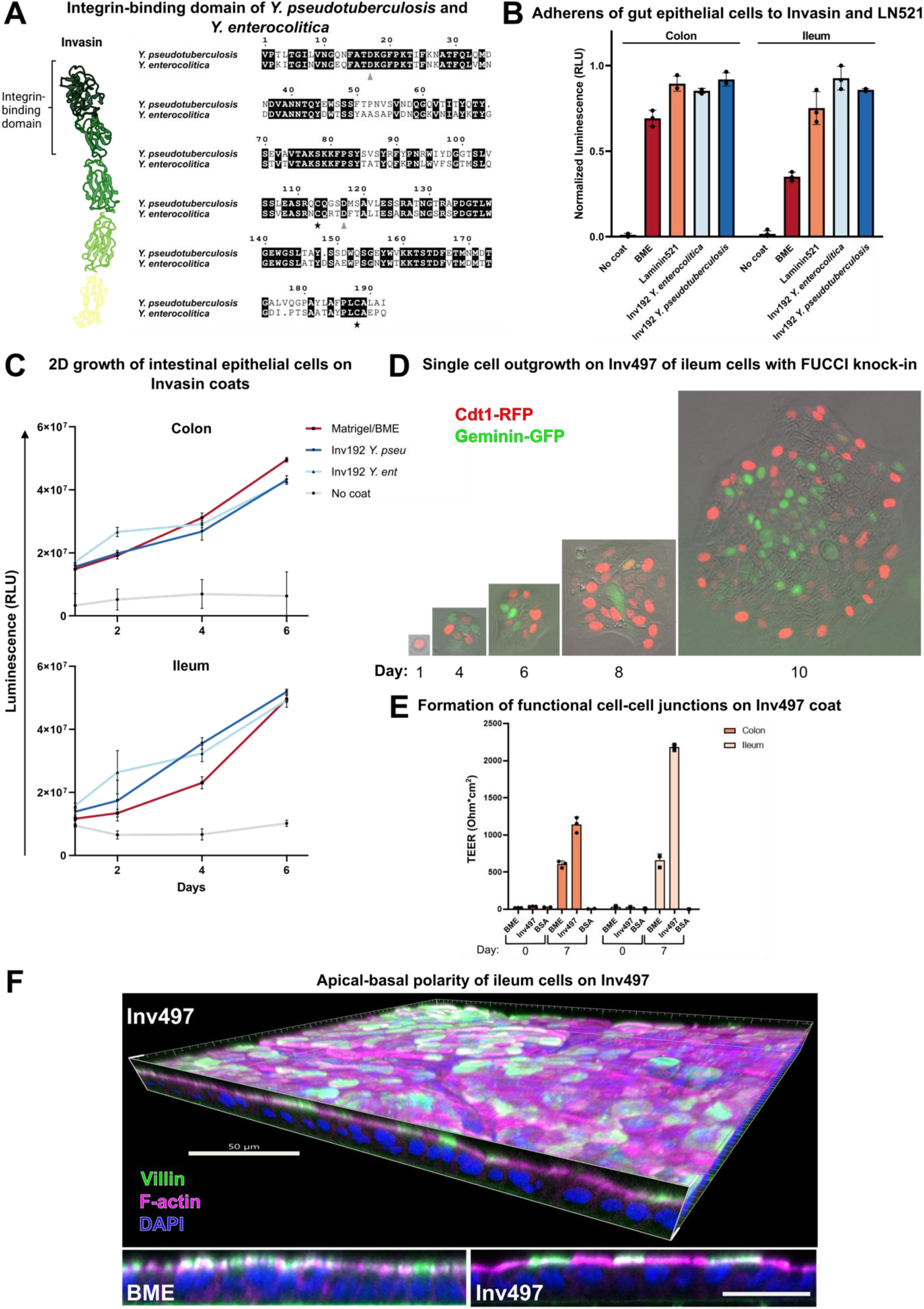
The integrin-binding domain of *Yersinia* Invasin supports epithelial cell adhesion, growth, junction formation and polarity. **A)** Structure of *Y. pseudotuberculosis* Invasin, with its C-terminal integrin-binding domain (from^42^). Sequence alignment of the integrin-binding domain of Invasin from *Y. pseudotuberculosis* and *Y. enterocolitica*. The asterisks indicate the cysteines involved in essential disulphide loop^43^. Grey arrows indicate conserved crucial amino acid residues involved in integrin interaction^34,44^. Alignment generated with the ESPript 3.0 tool. **B)** Quantification of adherence of organoid-derived colon/ileum single cells to coated BME, recombinant laminin521 (LN521) or Inv192 fragments from *Y. pseudotuberculosis* and *enterocolitica*, using a cellular ATP-driven luminescent assay (CellTiter-GLO). Relative light unit (RLU). Mean ± SD, n=3 **C)** Quantification of 2D growth of BME-established colon and ileum organoids on different coated wells, all started from 50.000 cells/well. The coats are BME, Invasin from *Y. pseudotuberculosis* (Inv192 *Y. pseu*), *Y. enterocolitica* (Inv192 *Y. ent*) or no coat. Mean ± SD, n=8 **D)** Culture started from 1 single cell from ileum 3D-established organoids expressing FUCCI cell cycle construct, growing on Inv497 of Yersinia *pseudotuberculosis*. Geminin-GFP indicates S (DNA synthesis), G2 (Cell growth) and M (Cell division, mitosis) phases of the cell cycle, whereas Cdt1-RFP indicates the G1 (Cell growth) phase of the cell cycle. **E)** Transepithelial electrical resistance (TEER) measurement of colon and ileum cells grown on Inv497, BME, or BSA coats, measured on day 0 and day 7. Mean ± SD, n=3 **F)** Epithelial monolayers of human ileum cells were grown on BME or Inv497 for 10 days. Staining for apical marker F-actin, apical brush border marker Villin, and nuclear marker DAPI. Scale bar in the top panel is 50 μm and in the lower two panels 25 μm.

We produced C-terminal fragments (192 aa and 497 aa) of the two *Yersinia* species in *E.coli* either in isolation or fused to Maltose-Binding Protein (MBP). Human ileum and colon epithelial cells, derived from organoid lines, spontaneously adhered to culture plates coated with the Invasin fragments of both species, as they did to a control coat consisting of a recombinant form of the ‘adult’ epithelial basement membrane protein laminin521 (Fig. 2B). Adhesion of human ileum and colon epithelial cells was equal between Inv192 and Inv497 and was not affected by the presence of MBP (Suppl. Fig. 3). Of note, similar adhesion has previously been shown for T lymphocytes and Hep-2 cells^35,39^. Since we observed no clear functional differences between the integrin-binding domains of *Y.enterocolitica* and *Y.pseudotuberculosis*, we performed all subsequent studies with the *Y.pseudotuberculosis* protein. Additionally, the Inv192-and Inv497-BMP fusion proteins behaved equally in these initial experiments and since the latter fusion protein was more easily produced (3 mg/ml per 50 ml bacterial culture), we elected to use this (‘Inv497’) for all subsequent experiments.

Importantly, the adherent ileum and colon cells expanded when organoid medium was added to the wells (Fig. 2C), as quantified using an ATP-depended luminescent light assay. We next performed clonogenicity assays, a particularly high bar for culturing epithelial cells. When human ileum organoid-derived cells were seeded at one cell/well in a 96-culture plate, we observed an average plating efficiency of 38.5% on an Inv497 coat, compared to our positive control Matrigel/BME-coat at 45.8%, while no clones grew out under ‘no coat’ or bovine serum albumin (BSA)-coat conditions (n=3, a total of 288 wells per condition was quantified). The outgrowth of a single ileum organoid cell, genetically modified with the Fluorescent-Ubiquitination-based Cell Cycle Indicator (FUCCI) reporter (from^40^), was followed over 10 days (Fig. 2D). Ileum epithelial cells grown on an Inv497 coat in transwell plates developed into a tight, confluent epithelial layer with functional cell-cell junctions as evidenced by the development of high levels of Trans-Epithelial Electrical Resistance (TEER), measured on day0 and day7 compared to a control protein coat BSA (Fig. 2E). Importantly, TEER levels were generally higher for sheets grown on Inv497 coats *versus* Matrigel/BME coats. Ileum organoid cells grown on an Inv497 coat displayed prominent apical-basal polarity, as observed with F-actin and the mature enterocyte brush border marker villin (Fig. 2F) (see for additional polarity markers Suppl. Fig. 4).

We next addressed the universality of this 2D culturing approach. We thus found that Inv497 coats supported adhesion and 2D growth of: 1) human airway cells^44^ (both starting from 3D-organoid-derived cells or from fresh lung tissue, Suppl. Fig. 5), of: 2) primary mouse intestine^1^ (Suppl. Fig. 6), of: 3) mouse lacrimal gland organoids^45^ (Suppl. Fig. 7) and of: 4) snake venom gland epithelial cells^46^ (Suppl. Fig. 8), all in the presence of the appropriate growth factor cocktails. Growth of epithelial 2D sheets was documented by Ki67 staining, an ATP-sensitive luminescent assay, and/or cell counting (Suppl. Fig. 4-8). Again, TEER measurements confirmed the formation of confluent epithelial layers of human airway, mouse intestinal, mouse lacrimal gland, and snake venom gland cells (see suppl. Fig. 5-8).

We then asked if an Inv497 coat can sustain long-term growth of primary human epithelial cells? After 1-2 weeks of culturing on Inv497-coated trans-well plates, epithelial sheets were enzymatically dissociated and replated on freshly coated inserts to form new 2D sheets. This procedure was repeated multiple times for human ileum, colon and airway epithelial cells, all derived from organoid lines previously established in 3D Matrigel/BME (see Fig. 3A). At the time of writing, 2D cultures of human ileum and airway had reached passage 24 (n=2, weekly passaged at a 1:2 ratio) and 14 (N=2, weekly passaged at a 1:3 ratio), respectively. Comparable results were obtained for mouse lacrimal gland cells^45^ and for snake venom gland epithelium^46^, all starting from organoids that had been established in 3D (see Suppl. Fig 7 and 8).

**Fig. 3:**
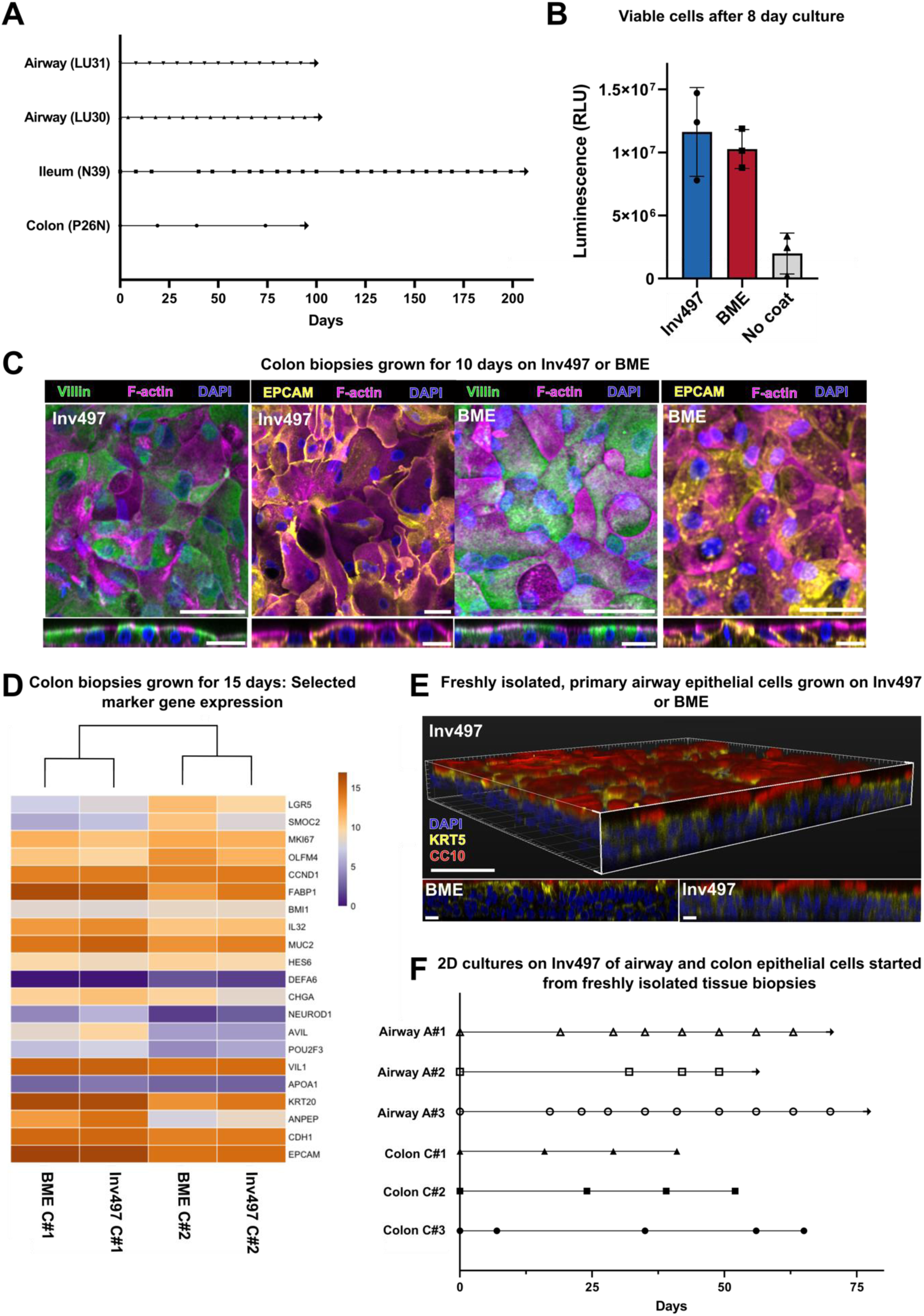
Long-term growth of epithelial cells. Establishment of cultures from primary tissue. **A)** Multiple passages in 2D on Inv497 of airway (LU30 and LU31^45^), ileum (N39^48^), colon (P26N^49^) 3D-established organoids. Dots represent passages. Passage ratio 1:2 for intestine and 1:3 for airway epithelial cells. **B)** Biopsies of healthy colon tissue cultured on BME, Inv497 or no coat. After 10 days, numbers of viable cells were quantified using CellTiter-GLO. Mean ± SD, n=3 **C)** Colon cells from tissue biopsies freshly grown for 10 days on BME and Inv497. Epithelial 2D sheets were stained for brush border marker Villin (green), pan-epithelial marker EPCAM (yellow), F-actin (purple) and nuclear marker DAPI (blue). Scale bar top panel: 50 μm and lower panels: 25 μm. **D)** Bulk RNA sequencing of colon biopsies from two different donors (lines C#1 and C#2) grown on BME or Inv497 for 10 days. Heat maps indicate selected markers: stem cells: LGR5, SMOC2 and OLFM4; cell cycle: CCND1, BMI1 and MKI67; enterocytes: ANPEP, FABP1, Il32, VIL1, APOA1, KRT20; goblet cells: MUC2; enteroendocrine cells: CHGA and NeuroD1, Paneth cells: DEFA6; Tuft cells: AVIL, POU2F3; pan-epithelial markers: CDH1, EPCAM. **E)** Freshly isolated, primary airway epithelial cells grown on Inv497 or BME, stained for basal cells (KRT5, yellow); club cells (CC10, red); and nuclear marker DAPI (blue). Scale bar top panel: 50 μm; lower panels: 25 μm. **F)** Multiple passages in 2D on Inv497 of primary epithelial cells from healthy colon biopsies from three different donors (lines C#1, C#2, C#3) and airway biopsies from three different donors (lines A#1, A#2, A#3). Dots represent passages. Weekly passage ratio for airway epithelial sheets was 1:3 and for colon 1:2.

Next, we tested the ability to start cultures from freshly isolated colon and airway biopsies. Colon tissue from three different donors was dissociated and cultured on Inv497 or on Matrigel/BME coats. Both coats were essential for outgrowth (Fig. 3B). Immunofluorescent staining for the epithelial marker EPCAM and the intestinal marker Villin identified the growing structures as intestinal epithelium (see Fig. 3C). Freshly isolated primary colon samples cultured on Inv497 or Matrigel/BME for 10 days were highly similar and contained all major cell types, as determined by bulk mRNA sequencing (Fig. 3D). Inv497 and Matrigel/BME 2D-cultures started from airway biopsies were characterized by immunofluorescent staining after 3 passages, and yielded identical pseudostratified epithelial sheets as visualized by staining for basal cells (Krt5) and club cells (CC10) (see Fig. 3E). Primary cultures from colon (N=3) and airway (N=3) biopsies could be expanded for at least four passages (see Fig. 3F), with the airway tissue at passage 10 at time of submission (weekly passage with a 1:3 ratio).

We then asked if long-term growth on an Inv497 coat still allows the generation of fully differentiated epithelial cells? We previously described a 3D human ileum epithelial organoid line (N39), genetically modified with triple fluorescent reporters, marking goblet (MUC2), enteroendocrine (CHGA) and Paneth (DEFA5) cells^49^. This line was maintained on a 2D Inv497-coat for 12 passages, after which the epithelial sheet was induced to differentiate as described previously^49^. The generation of goblet, Paneth and enteroendocrine cells appeared comparable between cells cultured in 2D on Inv497 or on Matrigel/BME (see Fig. 4A; and for a FACS-quantification Suppl. Fig.9). Electron microscopy supported the maturity of the cells (Fig. 4B).

**Fig. 4:**
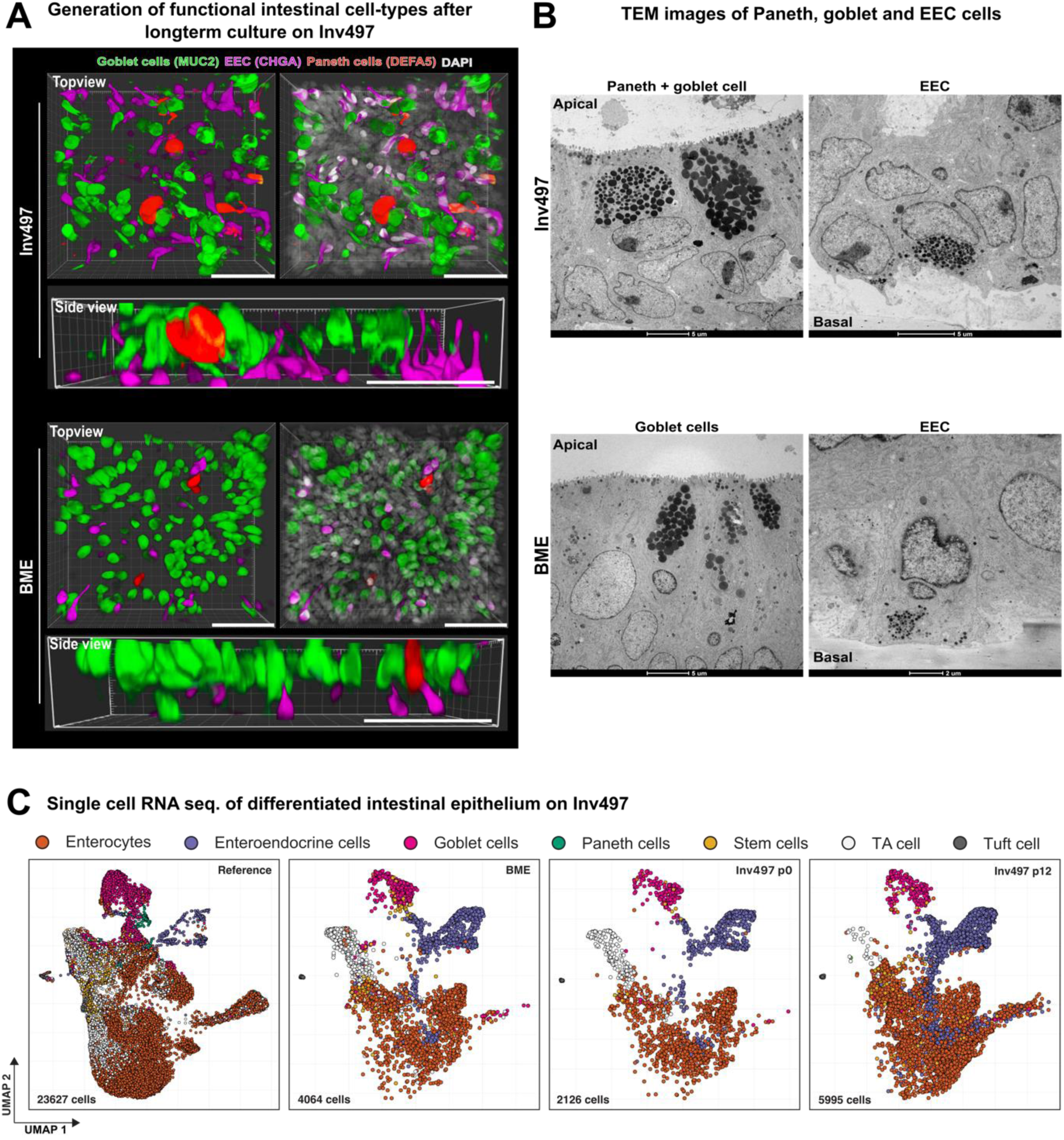
Long-term Inv497-culturing retains differentiation capacity to mature intestinal cell types. **A)** Top view and side view of differentiated ileum cells (triple knock-in reporter line^50^) after 12 passages on Inv497 or BME. Ileum epithelial cells differentiated towards Goblet cells (MUC2, green), Enteroendocrine cells (CHGA, purple), and Paneth cells (DEFA5, organge). Cell nuclei marked by DAPI (white). Scale bar is 50 μm **B)** Transmission electron microscopy (TEM) images of goblet cells, enteroendocrine cells (EEC) and Paneth cells upon differentiation after 12 passages on Inv497 or BME. Scalebar is indicated. **C)** UMAP of single cell RNA-sequence analysis of reporter ileum cells, differentiated after 12 passages on BME, early (P0) and late passage (P12) on Inv497. Left panel: a reference dataset from primary human ileum cells^51^. Descriptive cluster labels are given above the panels.

We then performed single cell RNA (scRNA) sequencing on these ileal cells differentiated after twelve passages (P12) or directly after plating (P0) on an Inv497 coat. We also sequenced organoid cells that were differentiated after 12 passages on a BME coat. After filtering, we captured 4064, 2125 and 5995 single cells from cells differentiated on BME, or on early and late passage on Inv497, respectively. We compared the transcriptomes of these cells to a reference dataset comprised of 23627 primary human ileum cells^50^. CCA integration identified shared markers between our dataset and the reference data. This allowed us to overlay the different datasets on a single UMAP projection (Fig. 4C), showing an overlap with fresh enterocytes, stem-, goblet-, transit amplifying-(TA), enteroendocrine-(EEC) and tuft cells. The number of Paneth cells captured in our dataset was too low to form a separate cluster; these cells were annotated manually based on known markers (DEFA5, PLA2G2A, REG3A). The number of TA cells and enterocytes varied between the different conditions: P12 Inv497 showed more fully differentiated enterocytes at the cost of TA cell numbers. Overall, this experiment indicated that the cells remained transcriptionally remarkably similar over multiple passages when cultured on Inv497. Comparison to primary ileal epithelial cells showed the presence of all major cell types.

Comparable results were obtained with human airway epithelial cells: epithelial 2D sheets directly derived from biopsies of 2 donors were passaged five times on Inv497 or BME coats, and were found to readily generate differentiated ciliated cells (acetylated tubulin) and Club cells (CC10) (see Fig. 5A) (Suppl. Fig. 10). The differentiated airway cells were also analysed by electron microscopy (see Fig. 5B), revealing a pseudostratified epithelium with prominent ciliated cells.

**Fig. 5:**
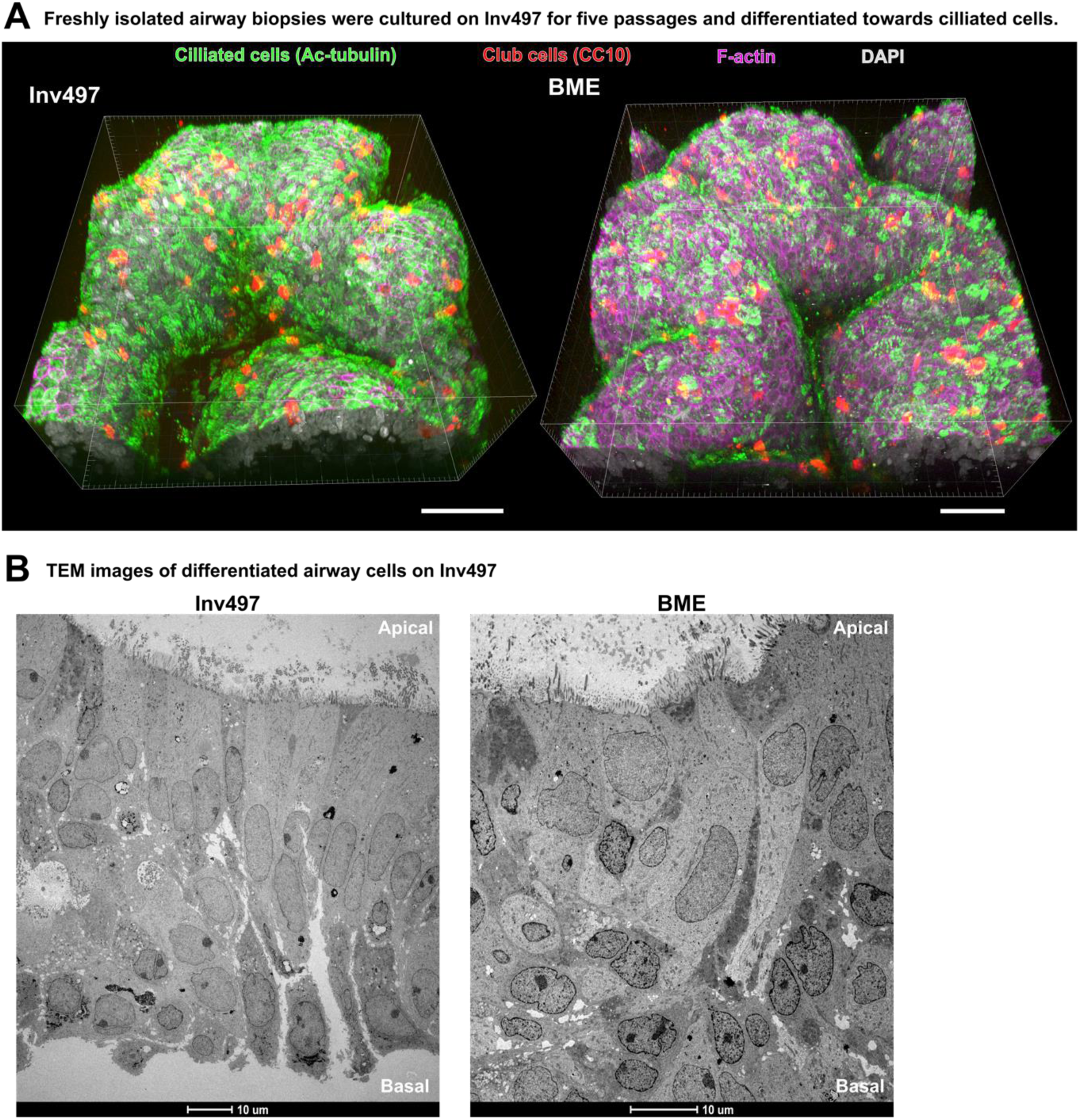
Long-term Inv497-culturing of freshly isolated airway biopsies retains differentiation capacity to mature cell types. **A)** Top view of Inv497 and BME airway cultures, started from freshly isolated biopsies (line A#1) and differentiated after five passages. Staining: ciliated cell marker: acetylated tubulin (green), club cell marker CC10 (red), F-actin (purple) and DAPI (blue). Scale bar is 50 μm. **B)** Transmission electron microscopy images (TEM) of differentiated airway cells cultured on Inv497 or BME showing a polarized pseudostratified epithelium with prominent ciliated cells.

## Discussion

Here we show that the truncated recombinant *Yersinia* protein Invasin, which uniquely activates multiple heterodimeric integrin receptors, can be used as an ECM-like ligand. Since most technologies for functional analysis of epithelia involve 2D culturing approaches, the “organoid sheet” format holds practical advantages over the 3D ‘in gel’ format: Basal as well as apical domains are directly accessible for infection-, uptake-or transport-studies, while functional integrity of junctional barriers can readily be assessed. Image analyses and high-throughput screening approaches are much simpler when executed in 2D.

A central aspect of organoid technology is the *in vitro* generation of multiple mature cell types, capturing key functions of the epithelium of interest^51^. Multiple mechano-physical studies have documented effects of 3D shape and substrate stiffness on the generation and maturation of such primary epithelial cells, be it as sheets or in the form of 3D organoids^52–55^. It was therefore surprising for us to note that -at least for the epithelia studied in detail-the diversity and maturity of the various epithelial cell types was maintained over multiple passages on Invasin-coated, high-stiffness 2D culture dish surfaces.

Much is known about the intracellular signals (and the central role of Focal Adhesion Kinase and Rho-type proteins) that emanate from the interaction of Invasin with integrin complexes. It will be of interest to document the roles of the individual signaling mechanisms in epithelial growth *in vitro*, yet we feel this to fall outside the scope of this initial study. As a next step, Invasin can be coupled to synthetic hydrogels to allow 3D culturing and variation of mechanical matrix properties.

While we realize that many applications remain to be explored and optimized, we believe that Invasin promises to represent a fully defined, cheap, versatile and animal-free alternative to Matrigel/BME.

## Material & Methods

### Organoid culture

Organoid lines were obtained as mentioned below (*organoid lines*) and maintained in corresponding media (see table 1) and Basement Membrane Extract (BME, R&D systems) BME^®^-domes. Organoids were passaged by mechanical disruption after 7 days (colon, ileum, snake, mouse lacrimal gland organoids) or 10-14 days (airway organoids). Passaging was done by aspirating medium and adding ice-cold DMEM/F12 (Gibco) on top of the BME^®^-domes. BME^®^ (R&D systems) droplets were disrupted by a P1000 pipette tip and resuspended before transferring into a 15-ml tube. The sample was centrifuged at 500g for 5 min, at 4°C. The DMEM/F12 (Gibco) medium was aspirated, and organoids were resuspended with P1000 (20x) and subsequently with P200 (20x) in fresh and cold DMEM/F12. Next, organoids were centrifuged, 500g for 5 min, at 4°C. The pellet containing the organoid cells were resuspended in cold BME^®^ and plated in +/-35 µl droplets of BME^®^ in suspension plates (Greiner, 657185). Plates were placed at 37°C (except for snake venom glands, incubation and culture was done at 32°C) for 10 minutes to solidify the BME^®^ droplets. Next, solidified BME^®^-domes were covered with corresponding epithelial expansion medium (see table 1).

**Table 1:** Table presenting media composition of different organoid types.

| Reagent | Source | Conc. | Colon<br>Ileum | Airway | Mouse<br>lacrimal<br>gland | Snake<br>Venom<br>gland | Mouse<br>Intestine |
| --- | --- | --- | --- | --- | --- | --- | --- |
| F12<br>advanced | Gibco | NA | + | + | + | + | + |
| Hepes | Gibco | 10 nM | + | + | + | + | + |
| Glutamax | Gibco | 1:100 | + | + | + | + | + |
| N-acetylcysteine<br>(Nac) | Sigma-Aldrich | 1,25<br>mM | + | + | + | + | + |
| Nicotinamide<br>(Nic) | Sigma-Aldrich | 10<br>mM | + | + (5 mM) | + | + |  |
| B27 | Sigma-<br>Aldrich | 1:50 | + | + (B27 +<br>vitA,) | +(B27+<br>vitA) | + | + |
| p38 inhibitor<br>SB202190 | Sigma-Aldrich | 10 µM | + | + (1 µM) |  |  |  |
| A83-01<br>inhibitor<br>TGF-β type I<br>receptor | Tocris | 0,5<br>µM | + | + | + | + |  |
| PGE2 | Tocris | 1 µM | + |  | + | + |  |
| Gastrin | Tocris | 10 nM |  |  |  | + (100<br>nM) |  |
| TGFα | Peprotech | 20ng/<br>ml |  |  |  |  |  |
| EGF | Peprotech | 50<br>ng/ml | + |  |  | + | + |
| FGF10 | Peprotech | 0,1<br>µg/ml |  | + | + | + |  |
| hHGF | Peprotech | 25<br>ng/ml |  |  |  |  |  |
| FSK | Peprotech | 1 nM |  |  | + |  |  |
| FGF7/KGF | Peprotech | 100<br>ng/ml |  | + (25<br>ng/ml) |  |  |  |
| CHIR | Sigma | 3 µM |  |  |  |  |  |
| NGS Wnt<br>Surrogate-Fc | #: N001<br>IpA | 0,5<br>nM | + |  |  |  | + |
| Rspodin3-Fc | #: R001<br>IpA | 3% | + | + (1%) | +(1%) | + (2%) | + |
| Noggin-Fc | #: N002<br>IpA | 2% | + | + (1%) | + (1%) | + | + |
| Penecillin<br>Streptomycin | Gibco | 100U/<br>ml | + | + |  | + | + |
| Primocin | Invivogen | 50<br>µg/ml | + | + |  | + |  |

### Organoid lines

- P26N colon organoids were previously described^48^.
- Human ileum organoids (N39) were previously described^47^.
- Airway organoids (LU30 and LU31) were previously described^44^.
- Snake venom gland organoids were previously described^46^.
- Mouse lacrimal gland organoids were previously described^45^.
- New lines from primary biopsies were processed as described^56^.

### Organoid media

#### Bacterial expression and purification of Invasin protein

An *E.coli* codon-optimized coding sequence of the C-terminal 192 amino acids of the *Inv* gene of *Yersinia pseudotuberculosis* and *Yersinia enterocolitica,* was synthesized by GeneArt (ThermoFisher) and cloned, using standard cloning protocols, into a pMalC2x vector (Addgene). An 8x-His tag, H3V-3C protease cleavage site, transglutaminase FXIII sequence and a FLAG-tag was introduced at the C-terminus of the MPB-tag (C-terminus of Inv could not be modified otherwise you lose proteins function). Sequence-verified pMalC2x Invasin192 vector were transformed into SHuffle T7 K12 Competent *E.coli* cells (New England Biolabs). Protein induction was carried out in bacteria grown in Terrific Broth (VWR) medium with 1mM IPTG (Sigma) when OD_600_ optical density reached 0.8 for 4 hours at 26°C. Bacterial cells were collected using centrifugation (10.000 rpm, 10 min.). The obtained bacterial pellets were resuspended in lysis buffer (50 mM HEPES, 200 mM NaCl (pH 7.4), 1mg/ml lysozyme (Thermofisher), 25 U/ml benzonase (Sigma), protease inhibitor cocktail (Roche) and subsequently sonicated (10 cycles; 10 seconds ON and 20 seconds OFF). Upon centrifugation for 20 minutes at 11.000 rpm the soluble fraction (upper phase) was collected and used for Ni-Nta (Qiagen) purification step based on the 8x-His tag presented in the protein. Further purification was performed by size exclusion chromatography using a Superdex S200 increase 10/300 GL size-exclusion column (Cytiva) equilibrated with buffer (50 mM HEPES, pH 7.4, 200 mM NaCl). The purified proteins were analysed under reducing and non-reducing conditions, on a 4-15% TGX precast SDS-PAGE (BioRad) gel and visualized using InstantBlue Coomassie Protein stain (Abcam). Protein concentration was calculated based on protein characteristics, extinction coefficient and Abs.0.1% found with the online ProtParamTool, combined with NanoDrop measuring absorbance. Similar procedure was performed for the 497 amino acids long C-terminal domain of Invasin. Purified proteins were incubated with HRV-3C (1 µl cleavage enzyme / 200 µg Invasin protein) (Pierce) O/N at 4°C. MBP_His and HRV-3C_His were removed using Ni-Nta beads (Qiagen) to obtain purified Inv192 and Inv497 proteins. Cleaved products were analysed on 4-15% TGX precast SDS-PAGE (BioRad) gel as described above. Due to inefficient cleavage, we used MBP fusion proteins for epithelial cell cultures.

#### Confocal imaging of epithelial cells

Confocal imaging of epithelial cells (from the colon, ileum, airway) were grown in 2D in a transwell system (Greiner 12-well transparent, 3 µm pore size) or on a glass plate (Greiner) coated O/N at 4°C with 2% BME^®^ (R&D systems) or 5 µg/ml recombinant Invasin protein. 2D-grown organoid sheets were fixed with 4% formaldehyde (pH 7.4) at room temperature (RT) for 30 minutes. Subsequently, the fixed cells were washed with PBS 0.1% Triton (PBST)(ThermoFisher), permeabilized, using PBS 1% Triton, for 30 minutes at RT, followed by blocking of the cells with 2% BSA (Sigma) in PBS for 60 min at RT. The fixed and permeabilized cells were incubated O/N at 4°C with the following primary antibodies with the indicated concentration (see table 2). Upon extensive washing with PBST, the binding of primary antibodies was visualized by 2 hours of incubation at RT with secondary antibodies (1:1000). Phalloidin fluor conjugates (see table 2, 1:1000) were used to directly visualize F-actin. DNA was stained with DAPI (ThermoFisher) or Hoechst (ThermoFisher). 2D-grown epithelial cells were imaged on a confocal SP8 (Leica) microscope or Stellaris microscope and analysed using ImageJ and Imaris software.

**Table 2:** Antibody list for confocal imaging.

| Antibody | Conc. in PBS |
| --- | --- |
| Villin (Santa Cruz, sc-58897) | 1:1000 |
| Phalloidin488 (Life technology, A12379) | 1:1000 |
| Phalloidin647 (Sigma, 65906) | 1:1000 |
| EPCAM (Biolegend, 324210) | 1:1000 |
| Acetylated tubulin (Santa Cruz, 6-11B-1, sc-23950) | 1:200 |
| Uteroglobin/CC10 (R&D, MAB4218) | 1:250 |
| Ki67 (Millipore, AB9260) | 1:200 |
| KRT5 (AF138, PRB 160P-100; 905501) | 1:2000 |
| AIIB2 (kind gift of A.Sonnenberg) (1 mg/ml) | 1:1000 |

#### Adhesion assay epithelial organoids

Epithelial cells, derived from colon, ileum, airway, mouse intestine, mouse lacrimal gland and snake venom gland were grown in BME to form organoids. The 3D BME^®^-grown organoids were removed from BME^®^ using 1:100 dispase (5U/ml) (Thermofisher) added to organoid medium for 30 minutes at 37°C (32°C for snake venom organoids). Subsequently, organoids were washed twice with cold DMEM/F12 and organoids were dissociated into single cells using TrypLE (Gibco) at 37 °C for 15 min, with repeated mechanical shearing using P1000 pipet. Single cells were washed twice in DMEM/F12 medium. 96-well plates (Corning) were coated overnight at 4°C with 5 ug/ml recombinant Invasin proteins or 2% BME (R&D systems) in cold PBS milliQ. After coating overnight at 4°C, protein solutions were removed, and the coated wells were blocked with 2% BSA (Sigma) in PBS for 30 min. at RT. Equal number of cells were incubated in DMEM/F12 for 45 minutes at 37°C (32°C for snake venom gland cells) to allow cell attachment. After washing the wells twice with DMEM/F12 medium, the number of adherent cells was quantified using CellTiterGlo (Promega) and the ATP-driven luminescence was measured using a tristar multimode plate reader (Berthold).

#### Colony forming assay on Invasin protein

A 96-well plate (Greiner) was coated O/N at 4°C with 5 µg/ml of Invasin protein or 2% BME^®^ diluted in PBS. Ileum N39 organoids that have been growing in BME^®^ droplets for 7-days, were treated with 1:10 (5U/ml in culture medium) Dispase (Thermofisher) at 37°C for 30 minutes, washed twice with cold DMEM/F12 medium, and dissociated to single cells using TrypLE (Gibco). DAPI-negative cells were FACS (BD, influx) sorted (1 cell/well) into the protein coated 96-well plates containing 100 µl intestinal organoid growth media supplemented with 10 µM Y-27 (Stem cell technologies). Growth of colonies were analysed using brightfield and fluorescent images with EVOS microscope. After 25 days the total number of colonies was counted in a blinded manner and later also quantified based on an ATP-sensitive luminescent assay (CellTiterGLO, Promega).

#### Growth assay of epithelial cells in 2D

Plastic 96-well plates (Greiner) were O/N coated at 4°C with 5 µg/ml Invasin protein in PBS or 2% BME^®^ (R&D system) diluted in PBS. Next day solution was removed, and epithelial organoids (3D-established colon, ileum, airway, mouse intestine, lacrimal gland, snake venom gland organoids) were released from the BME^®^ matrix with 10U/ml Dispase (Thermofisher), 30 minutes at 37°C (32°C for snake venom organoids). Subsequently, organoids were washed twice with cold DMEM/F12 and treated with TryplE (Gibco), 15 minutes at 37°C, with repeated mechanical shearing using P1000 to dissociate organoids to single cells (The dissociation of single cells was carefully monitored using a brightfield microscope). Single cells were washed twice with cold DMEM/F12 and the number of dissociated cells were counted with trypan blue (VWR) cell counting using an hemocytometer. Same number of cells (50.000 cells / well) per epithelial organoid line were loaded in the coated wells. Growth was quantified using CellTiterGLO on different days and luminescence was analysed on a Tristar multimode plate reader (Berthold). Same procedure as described above was performed to measure TEER over multiple days in transwell systems or quantify the number of cells growing on the Invasin coat or BME^®^. Quantification of the number of cells on different days was done using Trypan blue cell (VWR) counting and hemocytometer.

#### Passage epithelial 2D cultures

Organoids (Colon, ileum, airway, snake, mouse lacrimal gland) grown in BME^®^ matrix were treated with Dispase (10 U/ml) to remove all traces of BME^®^ components. After washing the Dispase-treated organoids with cold DMEM/F12, organoids were dissociated with TryplE to single cells (carefully analyse dissociation with brightfield microscope), this reaction was stopped by washing cells two times with DMEM/F12. Approximately 50.000 cells/well of epithelial cells were cultured on O/N coated transwell inserts (Greiner Bio-one; transwell systems 24-well inserts 3 µm pore size) with 5 µg/ml Invasin or 2% BME^®^ diluted in cold PBS (coating done as described above). Inserts/wells with growing epithelial cells were refreshed with 200 µl corresponding expansion medium every two days (transwell system medium was refreshed with 200 µl expansion medium in upper and 500 µl expansion medium in bottom compartment). When epithelial cells reached 100% confluency cells were passaged by adding pre-warmed 100 µl Dispase (10 U/ml) to expansion medium in the lower compartment and 10 ul to the upper compartment of the transwells for 10 minutes at 37°C. After 10 minutes, complete epithelial sheets were collected with P1000 pipet by pipetting up-and down (epithelial sheets can be seen in the pipet tip) and washed 2x with cold DMEM/F12. Washing was done using centrifugation steps of 500 rpm, 4°C for 1 minute. To check if all cells were collected, wells were analysed using a brightfield microscope (Thincerts and wells should be empty), collect remaining cells in transwells, if present, with 500 ul DMEM/F12. Epithelial sheets were enzymatically (TryplE, reaction was stopped with DMEM/F12) dissociated to smaller clumps of cells. Dissociation was carefully monitored using brightfield microscope. Cell clusters were pelleted down (400 rpm, 4°C for 5 minutes) and resuspended in epithelial-corresponding expansion media supplemented with 10 µM Y-27 and plated (passage ratio 1:2 (intestine) or 1:3 (airway, mouse lacrimal gland, snake venom gland) on new Invasin or BME^®^ coated wells. After one day wells were refreshed with expansion medium without Y-27.

#### Differentiation of triple reporter ileum organoid line

Ileum N39 triple reporter line (Goblet cells; MUC2-GFP, enteroendocrine (EEC) cells; CHGA-iRFP and Paneth cells: DEFA5-dsRED) obtained from^49^ were passaged on the integrin-binding domain of Invasin (5 µg/ml) in transwell system with 3 µm pore size (Greiner Bio-one, 662630) using expansion medium. To analyse if small intestinal epithelial cells could retain the full functionality of an intestinal epithelium when growing on Invasin, epithelial cells were differentiated towards functional and well-known intestinal cell types like goblet, enteroendocrine and Paneth cells. This was done as described in^49^, with minor differences. The differences were as followed, Invasin-grown epithelial cells were cultured on transwell systems until 100% confluency. Differentiation was induced using a 2-step differentiation protocol. The first step is to add 500 µl of patterning medium to the lower compartment of the transwell system for 7 days (refresh medium every 2 days with patterning medium), the upper compartment is just air. The second step is to induce Paneth cells, 500 µl maturation medium (as described in^49^) was added to the lower compartment of the transwell system, with just air in the upper compartment. This culture condition was maintained for 7 days with refreshing maturation medium in the lower compartment every day. Differentiated cells were imaged with Leica stellaris8 STED microscope and analysed using Imaris software.

#### Differentiation airway 2D sheets

Freshly isolated airway tissue cells (two donors) or from the 3D-established organoids lines (LU30 and LU31) were cultured for at least five passages on the integrin-binding domain of Invasin (Inv497, 5 ug/ml) in transwell systems with 3 µm pore size (Greiner Bio-one, 662630) using expansion medium added to the upper and lower compartment of the transwell system. When the epithelial sheets reached 100% confluency (determined based on eye and TEER) differentiation was induced by adding complete PneumaCult Air-liquid interface medium (stemcell technology, #05001), including Heparin solution (stemcell technology, #07980) and hydrocortisone solution (stemcell technology, #07925) to the lower compartment of the transwell system. The upper compartment remained air. Protocol and concentrations used for media were described by stemcell technologies. After 2-3 weeks of culturing epithelial cells with complete PneumCult air-liquid interface medium, with every 2-3 days changing medium, differentiated cells were imaged using confocal SP8 (Leica) microscope and analyzed using Imaris software.

#### RNA extraction and sequencing

Epithelial cells from primary colon biopsies grown on Invasin or BME^®^ coats were collected using Dispase, as described before, and washed with DMEM/F12 medium. RNA purification was performed with the RNeasy Mini kit with DNase treatment (Qiagen). RNA integrity was measured using the Agilent RNA 6000 pico kit and concentrations were determined with Qubit RNA HS analyzer (ThermoFisher). RNA integrity number values of RNA samples were between 8.5 and 10. For data analysed in figure1 we used the dataset used in published in^57^.

#### Single cell RNA sequencing of ileum triple reporter line

For single cell RNA sequencing the triple reporter ileum N39 line, late passage (passage 12) and early passage (passage 0) were used for differentiation. The ileum epithelial cells were differentiated as described before in transwell systems (Greiner 24-well inserts 3 um pore size). The ileum intestinal sheet was treated with TrypLE (Gibco) until cells round-up and start to detach from the well. Single cells were collected and washed with cold DMEM/F12 and resuspended in cold PBS 0.04% BSA (FACS buffer). DAPI-negative cells were sorted with the BDinflux. Cells were again counted using an hemocytometer and around 1000 cells/µl, total 40.000 cells were loaded to droplet-based scRNA-seq using the 10x genomics platform. Libraries were prepared using the 10x genomics chromium 3’ gene expression solution v3.1 and sequenced on a NovaSeq6000 (Illumina). Read counts were analysed using Seurat (v5). The full script is available at https://github.com/GJFvanSon/Hubrecht_clevers/invasin_sc. In short, read matrix was filtered, allowing a minimum of 1500 features per cell. Cells were then clustered using standard settings and annotated based on known marker gene expression like described in^49^. Public single cell expression data was loaded into seurat and integrated with our dataset using the seurat intergratelayers function with standard settings.

#### Primary tissue processing

Three different donors for primary human colon lines were derived by intestinal endoscopic biopsy from the colon. Three airway lines were derived from biopsies after surgery. The patient’s informed consent was obtained, and the study was approved by the ethics committee of the University Medical Center Utrecht. Healthy tissue was confirmed by the pathologist at the hospital. Epithelial cells of the mentioned organs were isolated as described previously^2^. Crypts and single cells were cultured in corresponding organ expansion medium, including 10 uM Y-27, on a BME^®^ coat or Invasin coat at 37°C and 5% CO_2_. Primary tissue was maintained in culture by refreshing every 2 days the expansion medium and passage was done as described before.

#### Trans-epithelial electrical resistance (TEER) measurement

3D-established organoid lines (colon, ileum, snake venom gland, mouse lacrimal gland) were removed from BME^®^ using 10U/ml dispase for 30 min. at 37°C (32°C for snake venom gland organoids). Dispase-treated organoids were washed twice with cold DMEM/F12 using centrifugation (500 rpm, 5 min, 4°C), to remove last traces of BME^®^. Subsequently, organoids were dissociated with TrypLE (Gibco) at 37°C for 15 min. Single cells were washed twice with cold DMEM/F12. Single cells were suspended in expansion medium, supplemented with 10 uM Y-27, and plated on a pre-coated (similar as described before, 5 µg/ml Invasin and 2% BME^®^, O/N, 4°C) transwell system (Greiner 12-well transparent, 3 µm pore size). TEER measurements were done using Millicell ERS-2 voltohmmeter, protein coated transwells without cells were used as control sample for TEER quantification.

#### ITGA6 and ITGAV CRISPR-Cas9 base editing

For knocking out integrin α6 and integrin αV the following sequence was ordered containing a universal priming part and the sgRNA (underlined) for base editing: ITGA6=<u>GGGTAGCCATCTTGATTAAT</u>**C**GGTGTTTCGTCCTTTCCACAAG; ITGAV= <u>TTGAAACCATCCTGGTCCAG</u>**C**GGTGTTTCGTCCTTTCCACAAG. The bold marked cytosine residue is included to obtain sufficient transcription from the U6 promoter. Using side-directed mutagenesis with Q5 high-fidelity polymerase the sequence primer was cloned into SpCas9-vector #47511(addgene) together with a universal 5’/phos/-GTTTTAGAGCTAGAAATAGCAAGTTAAAATAAGGC primer. Upon PCR clean-up, T4-ligase and Dpn1 were used to relegate and generate the plasmid, this was done according to manufactures protocol (NEB). DNA was transformed into DH5α bacterial cells, and plasmids were verified using sanger sequencing (Macrogen) with sequence primer GGGCAGGAAGAGGGCCTAT.

#### Organoid electroporation

Organoid electroporation was performed as described previously^58^ with some modifications. Wild-type colon were treated with 1.25% (vol/vol) DMSO in expansion medium, two days prior to electroporation. Electroporation was done with 10^6 cells with BTXpress solution containing 2.5 µg ITGA6 sgRNA plasmid, 5 µg Piggybac transposase, 5 µg Piggybac hygromycin cassette^59^, 7.5 µg C>T Base editor_eGFP plasmid (Addgene plasmid #112100). Single electroporated cells were cultured in BME^®^ droplets in expansion medium containing Y-27 (stem cell technologies). Five days post transfection, hygromycin 100 µg/µl (Invivogen) was added to previous mentioned expansion medium. Fourteen days after selection, surviving-hygromycin selection clones were individually picked and sequenced with the following primers: ITGA6_fwd GTTGGTTCACGGCTCTTTCCCC; ITGA6_rev CACCACACCCAGTTAATTT, ITGAV_fwd TGAGAAGTTTCTGTTCTTCTAG, ITGAV_rev CATGAGAGTCCATTATTGAAAT. DNA was obtained using ZYMO Research DNA isolation kit.

#### Growth assay integrin knockout colon organoids

Wildtype colon (P26N), ITGA6, and ITGAV knockout (KO) organoids were cultured in BME-domes as described before. ITGA6 KO organoids were constantly cultured in presence of 10 μM Y-27 (stem cell technologies). Medium was aspirated from the wells and cold DMEM/F12 was added on-top of the BME-domes. Organoids were collected by mechanically disrupting the BME-domes using P1000 pipette. After centrifugation and aspirating the supernatant, organoids were dissociated with TryplE to single cells, this reaction was stopped by washing cells 2x with DMEM/F12. Single cells were pelleted down (300 rpm, 5 min, 4°C) and resuspended in 1 mL cold DMEM/F12. Live cells were counted using trypan blue and a hemocytometer. In 15 µl BME droplets 7500 cells were cultured for every condition. After 10 min incubation of the plate at 37°C, to solidify the BME-domes, 500 µl expansion medium was added supplemented with 10 nM of the β1-specific inhibitory AIIB2 antibody (kind gift from Arnoud Sonnenberg) or 10 μM Y-27. After 7-days of culturing with corresponding media, brightfield images were taken with EVOS cell imaging microscope and cultures were treated with CellTiterGLO (promega) to quantified viable cells.

#### Transmission Electron Microscopy (TEM)

For Transmission EM, airway and ileum epithelial cell cultures were cultured on Invasin coated transwell system (Greiner 12-well translucent, 3 µm pore size). Ileum organoids were differentiated as described above. Airway organoids were differentiated using PneumaCult airway medium and protocols (from Stem cell technologies). Differentiated cells were fixed with fixative (1.5% gluteraldehyde (sigma) / 0.067 M cacodylate (sigma) buffered to pH 7.4 / 1% sucrose (sigma) and washed and stored in washing buffer (0.1 M cacodylate (pH 7.4) / 1% sucrose). Samples were cryoimmobilized using a Leica EM high-pressure freezer (HPM10) and stored in liquid nitrogen until further use. Over 3 days, samples were freeze-substituted at -90°C in anhydrous acetone containing 2% osmium tetroxide and 0.1% uranyl acetate at -90°C for 72 hours and warmed to room temperature with a gradient of +5°C per hour (EM AFS-2, Leica). Sections of the samples were analysed in a Tecnai Spirit T12 Electron microscope equipped with an Eagle OCD camera (Thermo Fisher Scientific).

## Acknowledgements

We thank Theodore Grenier, Antonella Dost, Katarina Balazova for processing primary tissue from colon and airway biopsies. We also thank Thanasis Margaritis for facilitating single cell sequencing. We thank Martina Celotti for preparing airway organoid medium. We thank Lulu Huang for a kind gift to share FUCCI reporter organoids. We thank Hubrecht FACS facility, especially Joost Korver. Authors thank Anneta Brousali, Jorieke Salij and Onno Kranenburg of the Utrecht Platform for Organoid technology (U-PORT; UMC Utrecht) for patient inclusion and tissue acquisition. Supported by the NWO Gravitation project Material Driven Regeneration (024.003.013 ZWK MDR) funded by the Ministry of Education, Culture and Science of the government of the Netherlands JW, WdL, DK, HC.

## Author contributions

J.A.P.M.W. and W.L. conceived the initial approach.; J.A.P.M.W. performed cell culture experiments, purified proteins together with F.R. and C.J.; J.A.P.M.W. and D.K. did fluorescence staining experiments; J.A.P.M.W. performed gene deletion; G.S. analysed single cell RNA sequence and bulk RNA sequence data; R.S. performed mouse, snake and airway cell culture experiments. J.A.P.M.W. and L.L. did differentiation and analysed single cell RNA sequence. S.L. performed and analysed FACS experiments. C.L.-I, W.v.d.W and P.J.P. performed transmission electron microscopy. R.I. provided reagents and reviewed the manuscript. J.A.P.M.W., W.L. and H.C. interpreted the results and wrote the manuscript.

