## Supplementary material for "Integrin-activating Yersinia protein Invasin sustains long-term expansion of primary epithelial cells as 2D organoid sheets": Suppl Fig 1

**Suppl. Fig. 1: Ligand specificity and Invasin binding of Integrin α/β1 pairs**

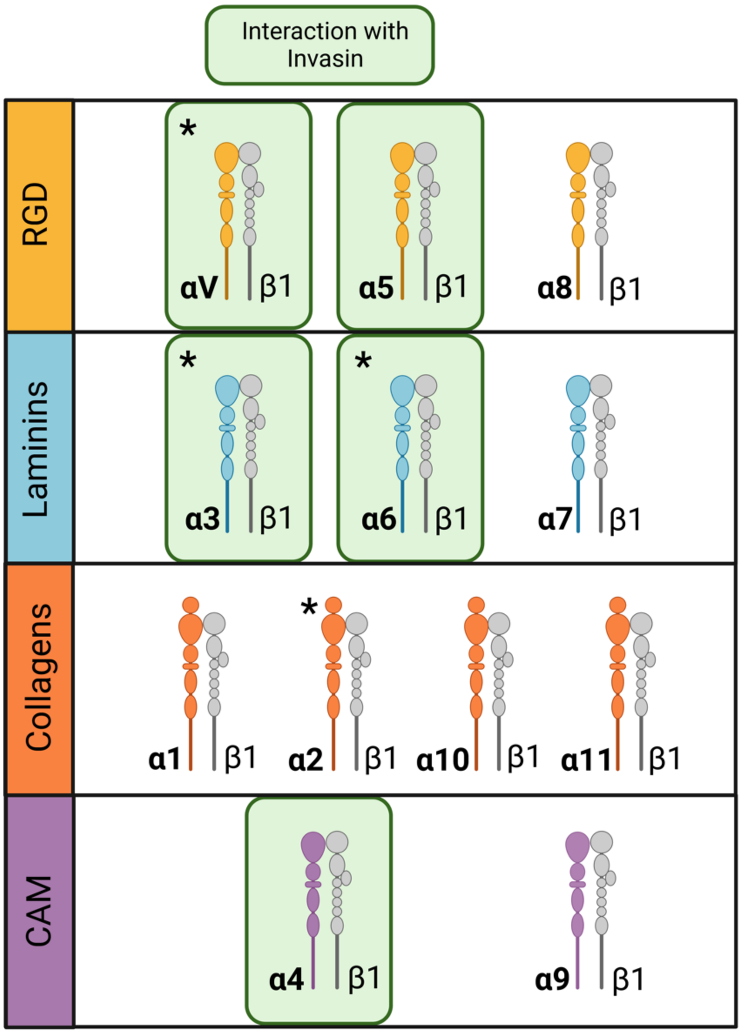


**Legend:** Integrin α/β1 pairs and their ligands. The universal integrin β1 (grey) associates with different integrin α subunits with unique ligand specificities: Collagens (orange), Laminins (blue), proteins containing the amino acid sequence RGD like fibronectin (yellow), or leucocyte-specific cell adhesion molecules (CAM; not expressed in intestinal epithelial cells) (purple). The integrin-binding domain of Invasin interacts with the heterodimers in the green boxes. Asterisks denote the integrins expressed in colon epithelial organoids (figure 1). Figure is modified from[8].
