## Supplementary material for "Integrin-activating Yersinia protein Invasin sustains long-term expansion of primary epithelial cells as 2D organoid sheets": Suppl Fig. 2

### Suppl. Fig. 2: ITGAV and ITGA6 CRISPR-Cas9 base editing of human colon organoids

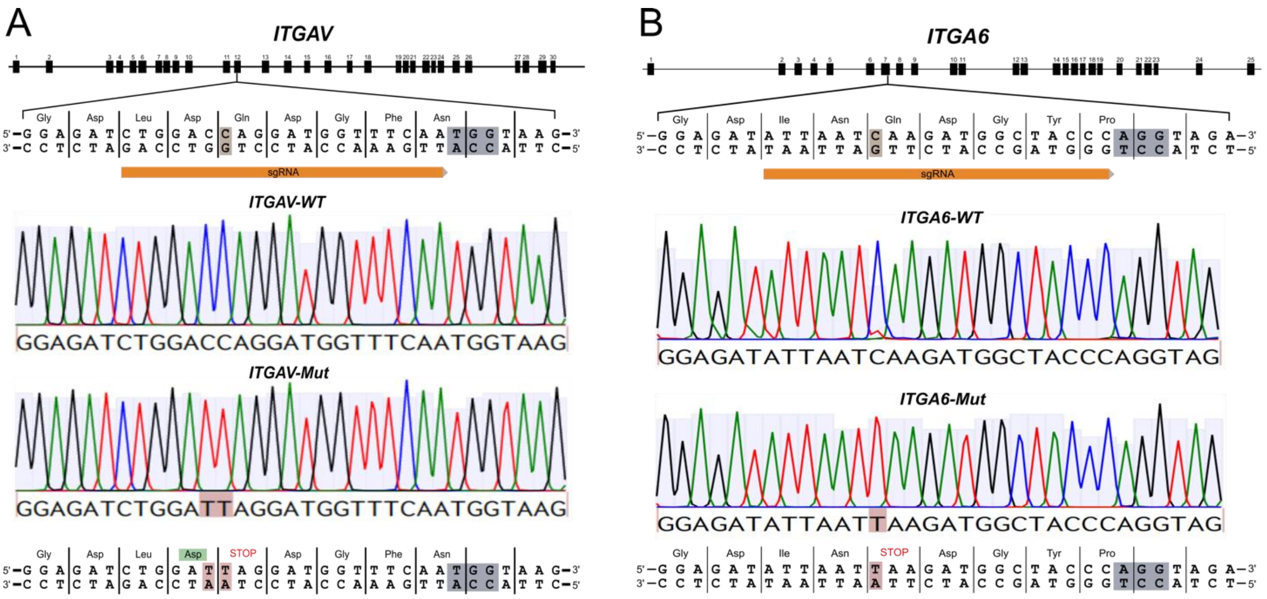


**Legend:** Sanger sequence validation of ITGAV (**A**) and ITGA6 (**B**) gene knockout using CRISPR-Cas9 base editing (base C>T editing. Editing transforms a Gln amino acid into a STOP codon. ITGAV and ITGA6 exons are illustrated and identify the location of the designed sgRNA (orange) and the mutation. The targeted base is highlighted in brown, and the PAM sequence is identified in grey. Single colonies were grown in the presence of Y-27 (Rho-kinase inhibitor). Image represents 1 of 3 verified clones. The additional C>T mutation in ITGAV for amino acid Asp (green amino acid) is a silent mutation.
