## Supplementary material for "Integrin-activating Yersinia protein Invasin sustains long-term expansion of primary epithelial cells as 2D organoid sheets": Suppl Fig. 3

**Suppl. Fig. 3. Gut epithelial cells adhere to Inv192 and Inv497 from *Y. pseudotuberculosis*; no effect of the MBP-fusion protein**

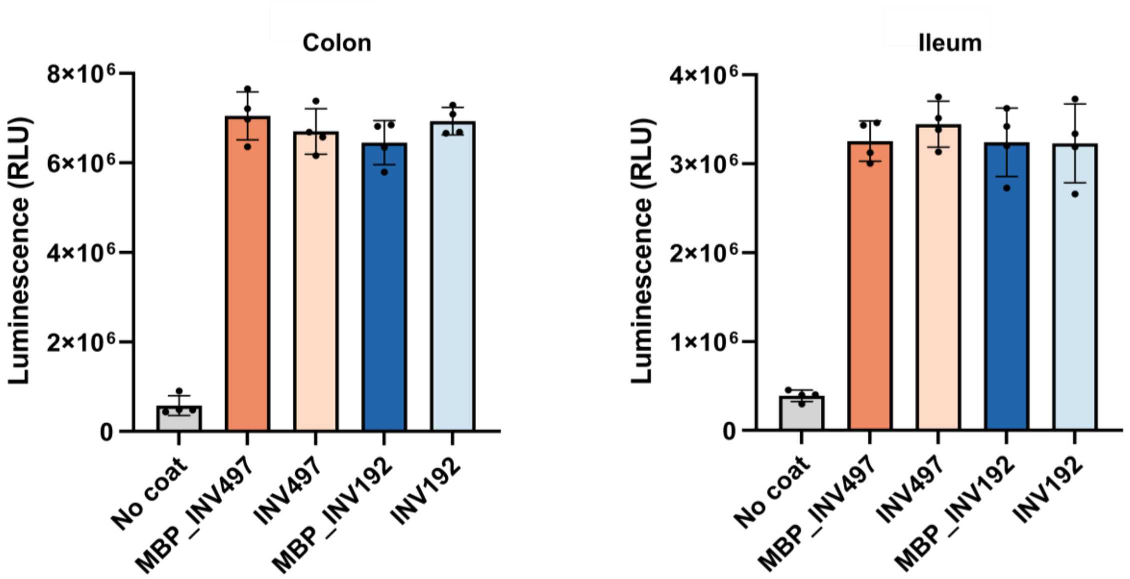


**Legend:** Quantification by CellTiter-GLO of colon and ileum organoid-derived single cells adhering to coats of Inv497 or Inv192, either as an isolated fragment or fused to Maltose binding protein (MBP). Identical protein concentrations (5 ug/ml)) where used for coating. RLU are given for four technical replicates for each condition. Means and standard deviation are indicated.
