## Supplementary material for "Integrin-activating Yersinia protein Invasin sustains long-term expansion of primary epithelial cells as 2D organoid sheets": Suppl Fig. 4

### Suppl. Fig. 4. Cell division and polarity of intestinal epithelium grown on Inv497 coats

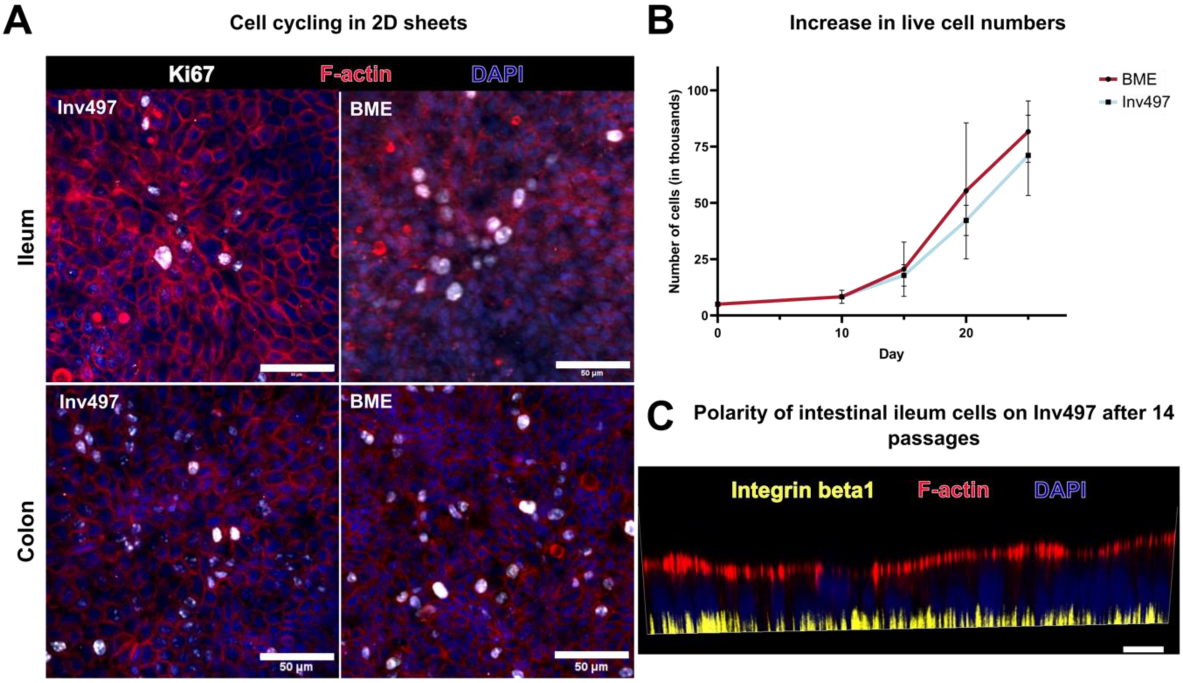


**Legends: A)** Top view of human colon and ileum cells grown in 2D on BME or Inv497 coats. Staining for proliferation marker Ki67 (white), F-actin (red) and DAPI. Scale bar is 50 $\mu$m**. B)** Human ileum organoid cells cultured for the indicated days; quantification of live cell numbers was done using trypan blue counting with a hemocytometer (n=3), values are in thousands. Mean and standard deviation are indicated. Of note, cells did not adhere and rapidly died on non-coated plates **C)** Human ileum 2D organoid sheets cultured on Inv497 for 14 passages maintain polarity. Basal marker integrin $\beta$1 (yellow), apical marker F-actin (red) and nuclear marker DAPI (blue). Scale bar is 25 $\mu$m.
