## Supplementary material for "Integrin-activating Yersinia protein Invasin sustains long-term expansion of primary epithelial cells as 2D organoid sheets": Suppl Fig. 5

**Suppl. Fig. 5 Human airway epithelial cells expand and maintain polarity upon long-term culture on Inv497 coats**

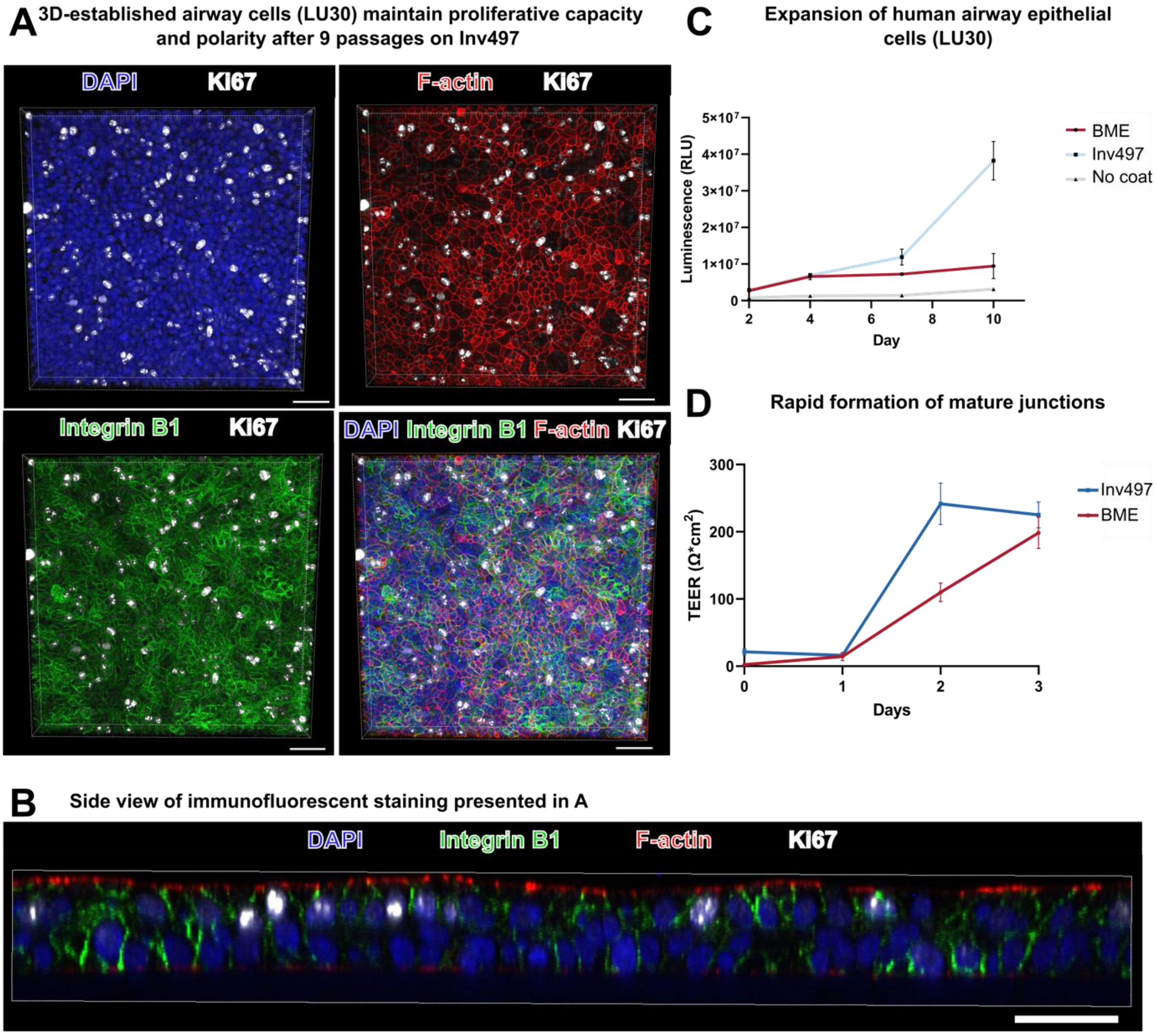


**Legends: A)** Airway epithelial organoid cells (LU30 from[45]) were cultured for 9 passages on Inv497 coat and stained for integrin β1 (green), proliferation marker Ki67 (white), apical marker F-actin (red) and DAPI (blue). Images represent top view. Scale bar is 50 μm. **B)** Side view of immunofluorescent staining presented in **A.** Scalebar is 50 μm. **C)** Human airway epithelial cells (LU30) were cultured on Inv497, BME or no coat. CellTiter-Glo was performed over multiple days. Standard deviation and mean are indicated of five technical replicates. **D)** Freshly isolated human airway cells were cultured for 3 passages on Inv497 and BME. After 3 passages, cells were replated, and TEER was measured over three days. Standard deviation and mean of three technical replicates are indicated.
