## Supplementary material for "Integrin-activating Yersinia protein Invasin sustains long-term expansion of primary epithelial cells as 2D organoid sheets": Suppl Fig. 6

### Suppl. Fig. 6 Cells directly harvested from mouse intestinal tissue grow on Inv497

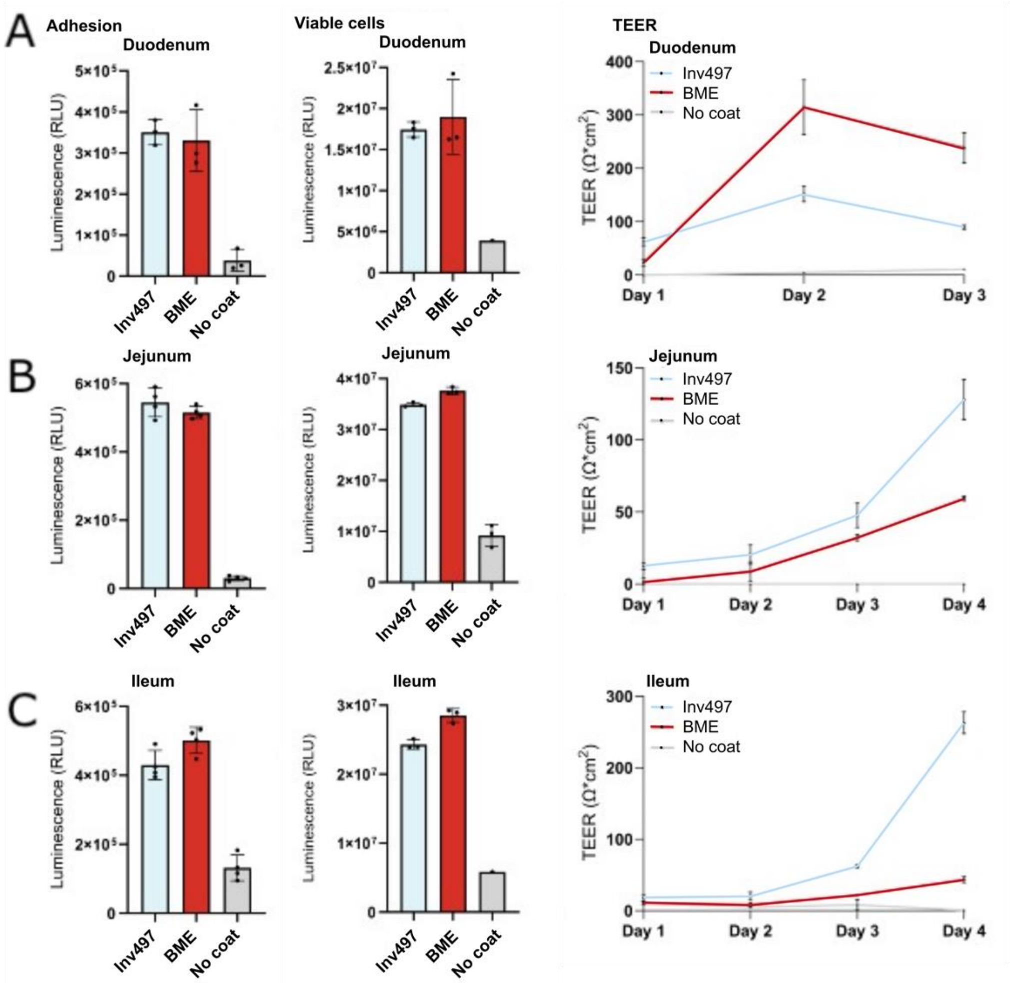


**Legends: A-C** Freshly isolated epithelial cells from mouse duodenum, jejunum and ileum adhere to Inv497 (left, first panel). Mean ± SD n=4. Viable cells were measured using CellTiterGLO after 4 days of culture on Invasin (Inv497), BME or no coat (middle panel). Mean ± SD, n=3, N=3. TEER increased rapidly over 4 days of culture. Mean ± SD n=4, N=3.
