## Supplementary material for "Integrin-activating Yersinia protein Invasin sustains long-term expansion of primary epithelial cells as 2D organoid sheets": Suppl Fig. 7

**Suppl. Fig. 7 Adhesion and growth of mouse lacrimal gland epithelial cells on Inv497**

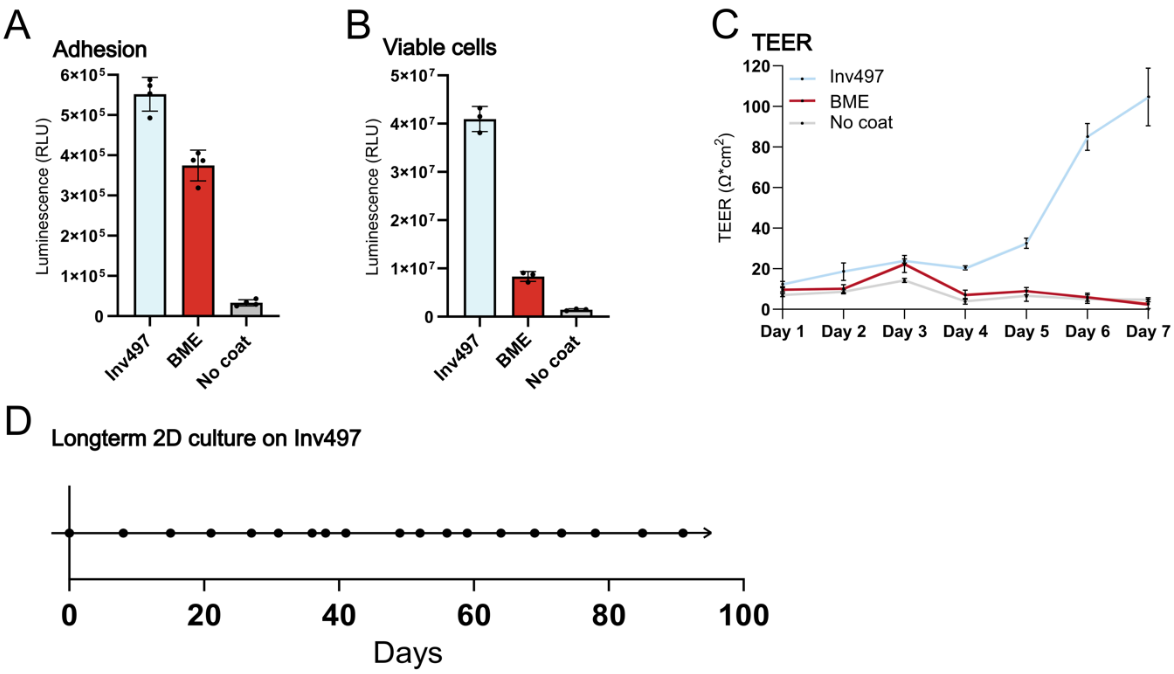


**Legends: A)** 50.000 cells mouse lacrimal gland organoid-derived single cells (from[46]) adhering to Inv497, BME or no coat. Assayed by CellTiterGLO. The mean of the relative light units (RLU) ± SD, are given for four technical replicates. **B)** Quantification of viable cells of mouse lacrimal gland after 7 days using CellTiterGLO. Mean ± SD, n=3. Started from 25.000 cells/well. **C)** TEER of mouse lacrimal gland cultures over time. Cultures were started with 25.000 cells/well. Means ± SD, n=3. **D)** Timeline of long-term culture of mouse lacrimal gland organoid-derived cells on Inv497, with every dot indicating a passage. The line could be passaged for at least 18 times and growth remained exponential. At the time of writing, cells were still in culture with a weekly 1:3 passage ratio.
