## Supplementary material for "Integrin-activating Yersinia protein Invasin sustains long-term expansion of primary epithelial cells as 2D organoid sheets": Suppl Fig. 8

**Suppl. Fig. 8: Organoid-derived snake venom gland epithelial cells grow on Inv497 coats**

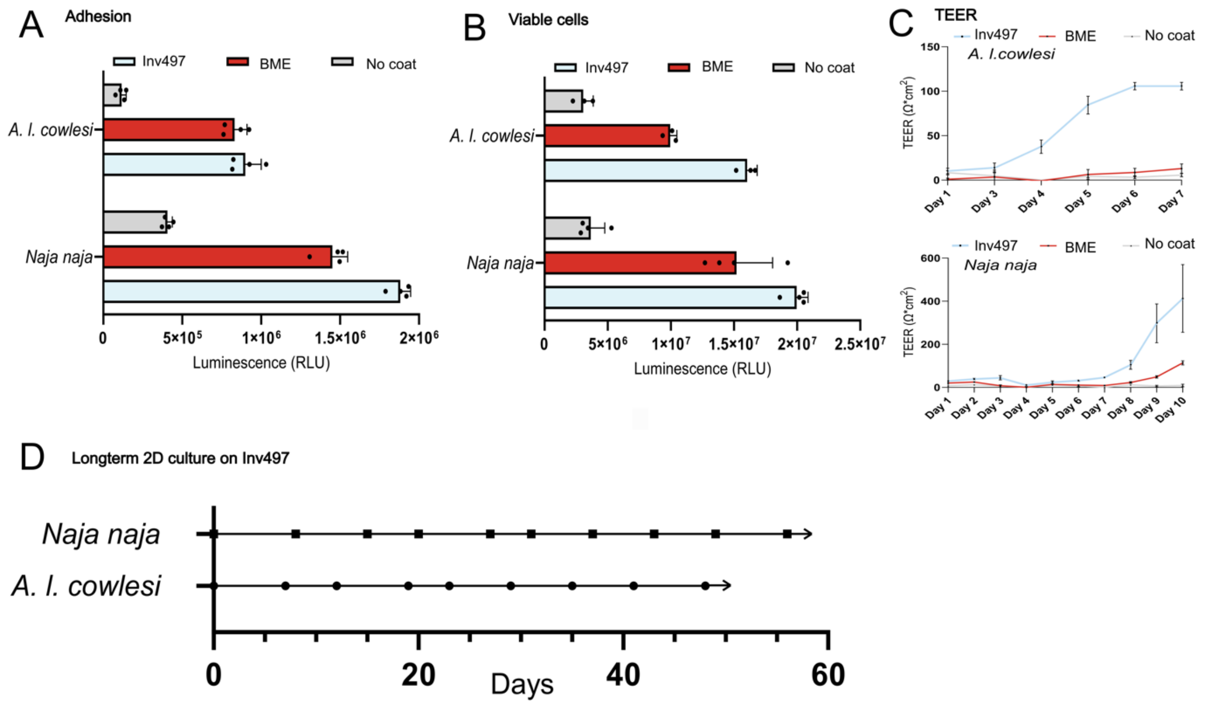


**Legends: A)** Snake epithelial cells (from[47]) recognize Invasin as extracellular matrix protein using an adhesion assay. Adhesion assay was performed with 30.000 cells/well. Figure represents quantification of *Aspidelaps lubricus cowles* (*A. l. cowlesi*) and *Naja naja* venom gland organoid-derived single cells adhering to a coat of Invasin (Inv497), of BME or no coat, using CellTiterGlo. Mean ± SD, n=4. **B)** Quantification of growth of *Aspidelaps lubricus cowlesi* (*A. l. cowlesi*) and *Naja naja* venom gland cells after seven days of culture on Inv497, BME or no coat. **C)** TEER of *A. l. cowlesi* (started with 35.000 cells/well) and *N. naja* (Started 20.000 cells/well) venom gland 2D cultures over time. Mean ± SD, n=3. **D)** Timeline of venom gland cells of the indicated snake species in culture on Inv497 coat, with each dot indicating a passage. At the time of writing, exponentially growing cells were still in culture with a weekly 1:3 passage ratio.
