## Supplementary material for "Integrin-activating Yersinia protein Invasin sustains long-term expansion of primary epithelial cells as 2D organoid sheets": Suppl Fig. 9

**Suppl. Fig. 9: FACS quantification of Paneth, goblet and enteroendocrine differentiation upon long-term culture of ileal cells on Inv497**

**A**

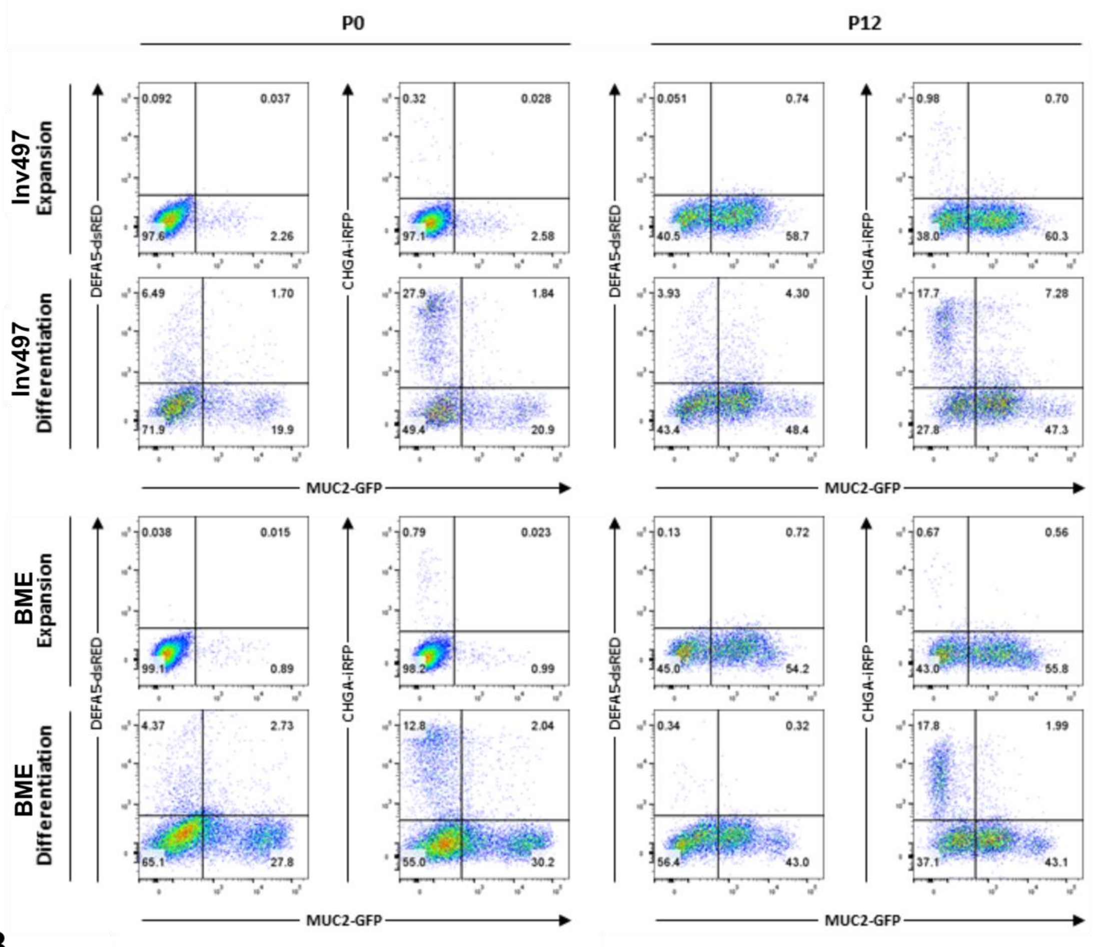


**B**

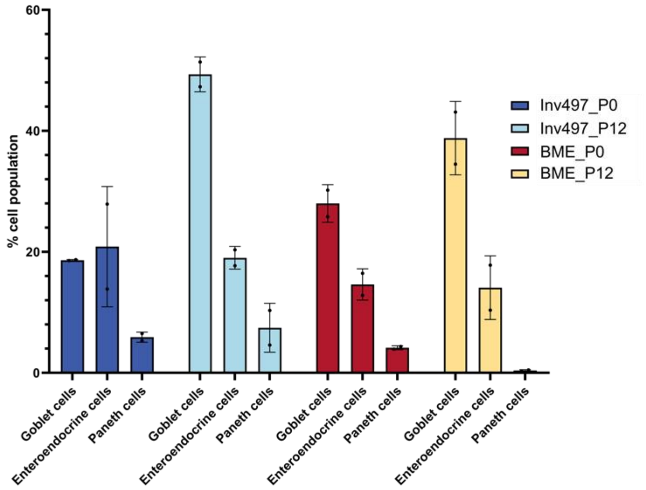


**Legends: A)** FACS analysis of human ileum triple reporter cells (from[50]) marked with MUC2-GFP (Goblet cells), CHGA-iRFP (Enteroendocrine cells), DEFA5-dsRED (Paneth cells), differentiated after 12 passages on Inv497 or BME. DAPI-negative cells were excluded; the remainder was analyzed for the expression of the indicated fluorophores. **B)** Quantification of cell type percentages (Goblet, Enteroendocrine, and Paneth cells) of two technical replicates of early passage (P0) and late passage (P12) ileum cells grown on Inv497 or BME. Mean and standard deviations are indicated.
