## Supplementary material for "Integrin-activating Yersinia protein Invasin sustains long-term expansion of primary epithelial cells as 2D organoid sheets": Suppl Fig. 10

**Suppl. Fig. 10 Freshly isolated airway tissues cultured on Inv497 maintains its potential to differentiate**

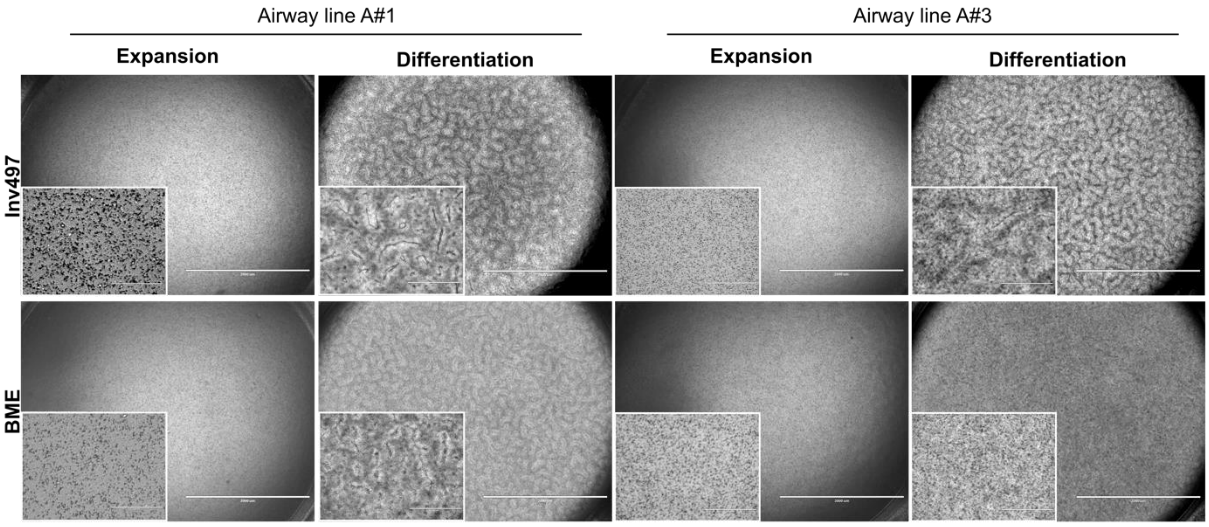


**Legend:** Brightfield images of primary airway epithelium (two donors: line A#1 and line A#3) at passage five on the c-terminal domain of Invasin (Inv497) and BME in expansion and differentiation medium. Scalebar is 2 mm.
